# In vitro exposure to non-antipseudomonal antibiotics (NAPA) induces *Pseudomonas aeruginosa* resistance to antipseudomonal antibiotics (APA)

**DOI:** 10.64898/2026.02.13.705873

**Authors:** David Lasry, Luke B. Harrison, Reggie Bamba, Rachel Corsini, Cedric P Yansouni, Matthew P. Cheng, Todd C. Lee, Alexander Lawandi

**Affiliations:** Division of Infectious Diseases, Department of Medicine, McGill University Health Centre, Montréal, Québec; Division of Medical Microbiology, Department of Laboratory Medicine, McGill University Health Centre, Montréal, Québec; Department of Critical Care Medicine, McGill University Health Centre, Montréal, Québec

**Author notes:** Corresponding author: Alexander Lawandi, MDCM, MSc, MHS, FRCPC, 1001 Boul Décarie, E05.1709, Montréal, Québec, H4A3J1. these authors contributed equally (David Lasry, Luke B. Harrison).

**Keywords:** *Pseudomonas aeruginosa*, antibiotic resistance, collateral resistance, efflux pumps, AmpC, directed evolution

## Abstract

**Background:** *Pseudomonas aeruginosa* readily evolves antimicrobial resistance through regulatory plasticity and stress-adaptive pathways. Clinically, antibiotics lacking intrinsic antipseudomonal activity are often favored with the assumption that they avoid selective pressure on *P. aeruginosa*. Whether subinhibitory exposure to such “non-antipseudomonal antibiotics” (NAPA) can nevertheless select for canonical resistance pathways remains incompletely defined. Methods: Three *P. aeruginosa* strains (ATCC 27853 and two bloodstream isolates) were serially passaged over 14 days in the presence of ertapenem, ceftriaxone, or moxifloxacin at one-third the baseline minimum inhibitory concentration (MIC). MICs for antipseudomonal antibiotics (meropenem, ceftazidime, ciprofloxacin) were measured at serial time points and after a 3-day antibiotic-free recovery (day 14). Whole-genome sequencing was performed longitudinally to identify mutations.

**Results:** NAPA exposure led to reproducible elevations in antipseudomonal MICs: ertapenem triggered up to a 29-fold increase in meropenem MIC, ceftriaxone up to a 31-fold rise in ceftazidime MIC, and moxifloxacin up to a 12-fold increase in ciprofloxacin MIC. Elevated MICs persisted on day 14 despite absence of further antibiotic pressure. Genomic analysis revealed convergent evolution of mutations in efflux regulator genes (*nfxB, nalC, nalD, amrR*) and the β-lactamase–regulating gene *dacB*, emerging during periods of MIC escalation and mapping to regulatory pathways governing efflux and AmpC expression.

**Conclusion:** Subinhibitory exposure to antibiotics without intrinsic antipseudomonal activity reproducibly selected for heritable multidrug-resistant phenotypes in *P. aeruginosa*. Resistance arose through convergent evolution in regulatory genes classically associated with direct antipseudomonal antibiotic pressure, demonstrating that resistance architectures can be selected independent of target engagement and underscoring the inevitability of collateral resistance under antibiotic stress.

**Importance:** *Pseudomonas aeruginosa* is a major cause of hospital-acquired infections and is well known for its ability to develop antibiotic resistance. Clinicians often assume that antibiotics without activity against *P. aeruginosa* do not meaningfully influence its resistance behavior and are therefore safe choices when this organism is not the primary target. Our study challenges this assumption. We show that low-level exposure to such antibiotics is associated with increased resistance to key antipseudomonal drugs, even after the initial antibiotic exposure ends. Rather than arising from a single stable genetic change, resistance was accompanied by shifting genetic alterations in regulatory pathways that control drug efflux and β-lactamase expression. These findings highlight how antibiotic exposure can broaden the evolutionary pathways available for resistance in unintended pathogens. Recognizing these indirect and population-level effects of antibiotic use may help inform more cautious antimicrobial prescribing strategies in clinical settings.

## Background

*Pseudomonas aeruginosa* (PsA) is a major pathogen responsible for a wide range of infections, particularly in nosocomial settings. It represents the 6^th^ most common bacteria isolated in hospitals^1^, accounting for 8.7% of bloodstream infections^2^, 21.8% of ventilator-associated pneumonia^3^, and 16% of nosocomial urinary tract infections^4^. Preferentially infecting patients with comorbidities and immunocompromising conditions, the 30-day mortality of PsA bloodstream infections is above 25%^5^.

Owing to its virulence factors and intrinsic resistance to antimicrobials, *P. aeruginosa* infections are considered difficult to treat. Drug-resistant *P. aeruginosa* is becoming more prevalent and poses significant challenges to clinicians and infection control practitioners. A US surveillance study of over 24,000 clinical isolates found resistance rates of 33% to fluoroquinolones, 25% to extended-spectrum cephalosporins, 25% to carbapenems, and 20% were multidrug resistant (MDR)^6^. Identifying risk factors associated with resistance is critical as it can inform empiric therapy in critically ill patients prior to susceptibility results becoming available. Length of hospital stay, underlying comorbidities, and prolonged courses of antipseudomonal antibiotics (APA) appear to be the most consistently predictive factors in most models^7–9^.

Several case-control studies have linked exposure to non-antipseudomonal antibiotics (NAPA) as an independent risk factor for the development of isolates resistant to APA. Cohen et al.^10^ found that patients who had received ertapenem during their hospital stay were more likely to later be colonized or infected with pan-resistant PsA (HR,1.9, 95% CI,1.01-3.4). Similarly, Coppry et al.^11^ showed that receipt of beta-lactams inactive against PsA was an independent risk factor for the subsequent isolation of carbapenem-resistant PsA (OR, 1.101, 95% CI, 1.010-1.201). These studies draw into question the widely accepted concept that NAPA “spare” or avoid the development of drug-resistant *P. aeruginosa*.

While observational studies implicate non-antipseudomonal antibiotics (NAPA) as risk factors for subsequent isolation of resistant *P. aeruginosa*, the biological mechanisms underlying this association remain poorly characterized. We hypothesized that subinhibitory exposure to antibiotics lacking direct antipseudomonal activity could nevertheless impose sufficient selective pressure to favor canonical resistance pathways in *P. aeruginosa*. To test this, we employed a directed evolution framework exposing multiple *P. aeruginosa* strains to subinhibitory concentrations of representative NAPA, coupled with longitudinal phenotypic and genomic analysis.

## Methods

### Directed Evolution Assay

An *in vitro* directed evolution protocol was established to evaluate adaptive responses of PsA to subinhibitory antibiotic exposure (Figure 1). Three PsA strains were studied: a reference strain (ATCC 27853) and two bloodstream isolates obtained from the McGill University Health Centre (MUHC) clinical microbiology laboratory (designated here as isolates 124 and 985). Each strain was subcultured twice from frozen stock onto blood agar and incubated overnight at 37 °C in ambient air. From these, 0.5 McFarland suspensions were prepared in saline for subsequent assays.

**Figure 1.**
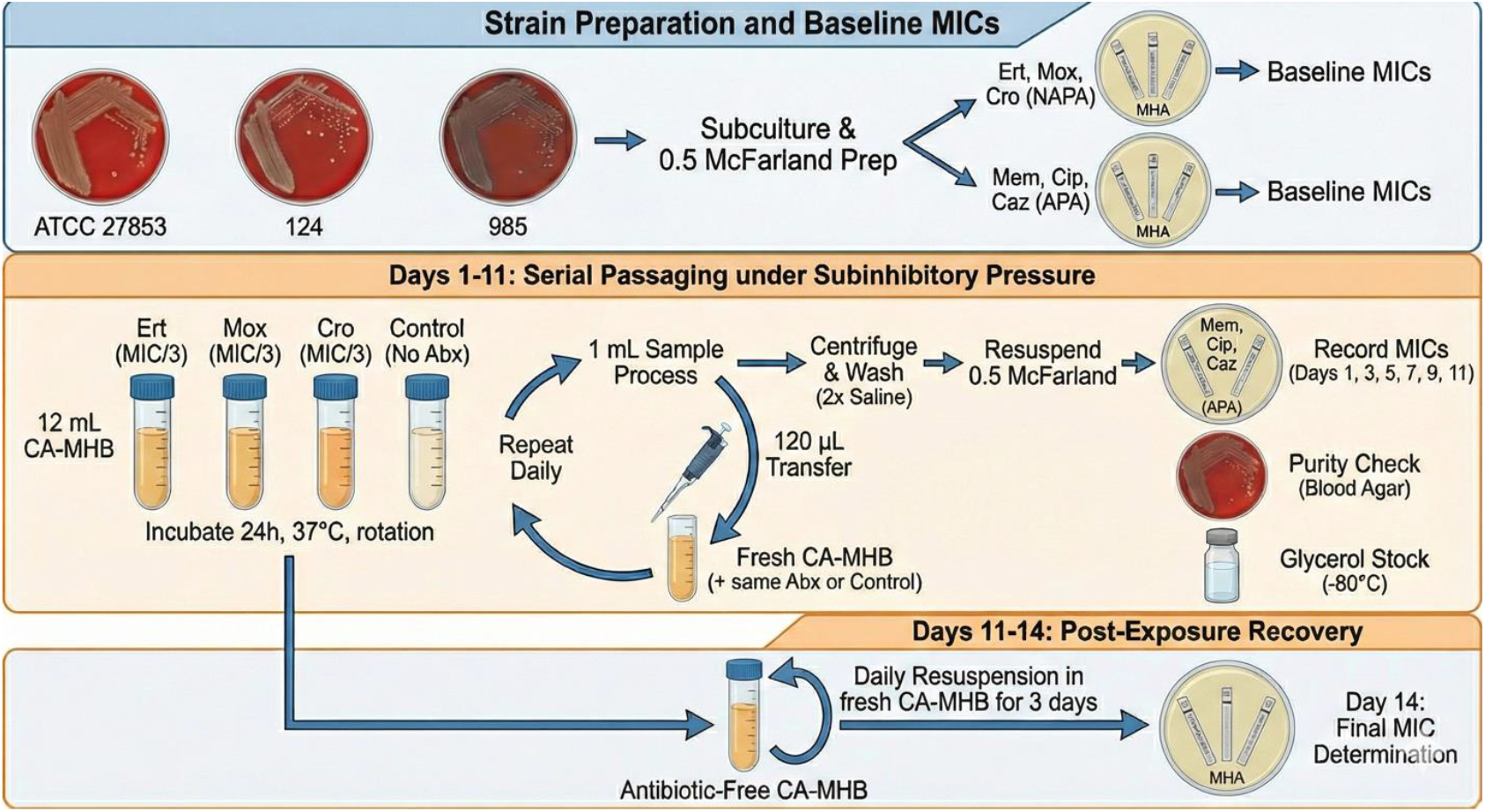
Adaptive Evolution Experimental Flow.

### Baseline Susceptibility Testing

Baseline minimum inhibitory concentrations (MICs) were determined using Epsilometer (E-test; bioMérieux, Marcy-l’Étoile, France) methodology on Mueller–Hinton agar (MHA). For each strain, two MHA plates were inoculated: one with ertapenem, moxifloxacin, and ceftriaxone (NAPA group), and the other with meropenem, ciprofloxacin, and ceftazidime (APA group). Plates were incubated at 37 °C for 18–24 h, and MICs were recorded according to Clinical and Laboratory Standards Institute (CLSI, M100, 34th ed) criteria.

### Culture Conditions and Antibiotic Exposure

For each strain, four 12 mL tubes of cation-adjusted Mueller–Hinton broth (CA-MHB) were prepared. Three tubes contained one of the NAPA antibiotics—ertapenem, moxifloxacin, or ceftriaxone—at one-third of the measured MIC (MIC/3) (as defined in Vinchhi at al.^12^), while the fourth served as an antibiotic-free control. Cultures were inoculated to a final density equivalent to 0.5 McFarland and incubated aerobically at 37 °C with constant rotation for 24 h.

### Serial Passaging and MIC Determination

Following incubation, 1 mL from each culture was centrifuged, and the bacterial pellet was washed twice with normal saline to remove residual antibiotic. The washed pellet was resuspended to a 0.5 McFarland density and plated onto MHA. E-tests for meropenem, ciprofloxacin, and ceftazidime were applied to determine MICs. Parallel inoculations were streaked onto blood agar to confirm purity, and 50 µL aliquots were preserved in 20% glycerol at –80 °C.

From each remaining liquid culture, 120 µL were transferred into fresh CA-MHB containing the same antibiotic concentrations (except for control cultures) to ensure continuous and uninterrupted exposure. Cultures were incubated for an additional 24 h under identical conditions. MICs were reassessed on days 1, 3, 5, 7, 9, and 11, with daily subculture into fresh antibiotic-containing medium.

### Post-Exposure Recovery Phase

After 11 days of serial passage, cultures were transferred into antibiotic-free CA-MHB and incubated for three additional days (with daily resuspension in freshly prepared CAMHB tubes) to evaluate persistence of MIC changes in the absence of drug pressure. Final MICs were determined on day 14.

### DNA Extraction and Whole Genome Sequencing

The frozen glycerol stocks of the three strains from days 0, 3, 5, 7, 11 and 14 were subcultured onto a blood-agar plate and grown overnight twice. When more than one colony morphology was present on a given BAP, a representative from each phenotypically distinct set of colonies was selected for sequencing (summary of samples and sequencing in Tables S1 and S3). Isolated colonies were suspended in MHB and sent to the McGill Genome Centre for DNA extraction and whole genome sequencing. The DNA extraction was performed using DNeasy 96 PowerSoil Pro QIAcube HT Kit (cat# 47021). The manufacturer’s protocol was followed without modification. Samples were prepared for sequencing using the Lucigen NextSeq AmpFree kit. Isolates were sequenced on 1 lane of NovaSeq 6000, Sprime PE150. Sequencing reads were retrieved, filtered and adapters were clipped using trimmomatic (v0.39)^13^ with the following parameters (2:30:10:2:True LEADING:3 TRAILING:3 SLIDINGWINDOW:5:20 MINLEN:50). Kraken2 with was used to verify the taxonomic identity of sequencing reads (Standard index, 2025/02/04). Baseline reference genomes for each strain were constructed as follows. For the ATCC 27853 strain, the corresponding reference genome and annotation was downloaded from the Pseudomonas genome database (https://www.pseudomonas.com/strain/show?id=15935, NCBI assembly GCF_016126955.1), then sequence reads from a sample at day 0 (sample 1_784603) were aligned against this genome using bwa-mem2^14^ and single nucleotide polymorphism (SNPs) called with GATK^15^, using the HaplotypeCaller and GenotypeGVCF tools. The reference genome was then updated with the called SNPs using the bcftools^16^ consensus tool to create an updated reference sequence and account for in-lab divergence from the reference genome assembly prior to experimental evolution. For the clinical strains, de novo assembly of the reads from each strain sequenced on day 0 (i.e. prior to experimental evolution, samples 23_784679_S23 and 45_784695_S39 for strains 124 and 985, respectively) was performed using spades (v4.0.0, option –isolate)^17^. Genome annotations for the de novo assembled reference genomes were created using PGAP (v2025-05-06.build7983)^20^.

Genomic changes in the serially sequenced samples were analyzed separately for each strain. SNPs were identified using a GATK workflow: first, raw sequencing reads were trimmed, filtered, and aligned to the respective reference genome, as described above. Then, the GATK tools HaplotypeCaller and GenotypeGVCF were used to perform joint genotyping for each of the three experiments. SNPs were then filtered using GATK’s VariantFiltration (options: --filter- expression “QD < 2.0” --filter-name “LowQD” --filter-expression “FS > 60.0” --filter-name “HighFS” --filter-expression “SOR > 3.0” --filter-name “HighSOR” --filter-expression “MQ < 40.0” --filter-name “LowMQ” --filter-expression “MQRankSum < −12.5” --filter-name “LowMQRankSum” --filter-expression “ReadPosRankSum < −8.0” --filter-name “LowReadPosRankSum”) and annotated using bcftools csq tool and the respective reference genome annotations. The resulting variant call files (VCFs) were manually inspected, and those mutations found at any allelic frequency in any control isolate or in areas of overall poor alignment or contig boundaries were removed (final SNP datasets in Tables S4-6).

## Results

### Change in MIC Over Time (Antibiotic-exposure vs Control)

Growth in media containing NAPA at a concentration of [MIC/3] conferred increased resistance to APA (Table 1). The inducible rise in MIC was most pronounced when measuring MICs to APA within the same class of the exposure antibiotic. Ertapenem conferred a maximum mean 29-fold increase in meropenem MIC; moxifloxacin conferred a 12-fold increase in ciprofloxacin MIC; ceftriaxone conferred a 31-fold increase in ceftazidime MIC. However, the induction of cross-resistance between antibiotics of different classes was also observed. For instance, both moxifloxacin and ceftriaxone exposure caused the meropenem MIC to rise appreciably, 6.5-fold, and 10-fold respectively. Moxifloxacin and ertapenem exposure did not consistently increase ceftazidime MICs in a significant way. Ertapenem exposure did not alter ciprofloxacin MICs appreciably either.

**Table 1.** Maximum fold-change in APA MIC relative to control after NAPA exposure. *Fold-change represents the maximum ΔMIC (intervention) / ΔMIC (control); day indicates when the maximum was observed during serial passage*.

| **NAPA exposure** | **APA MIC endpoint** | **PsA-ATCC** | **PsA-124** | **PsA-985** | **Mean** |
| --- | --- | --- | --- | --- | --- |
| Moxifloxacin | Meropenem | 8 (d5) | 4 (d9) | 7.9 (d5) | 6.5 |
|  | Ciprofloxacin | 10.5 (d9) | 3.9 (d11) | 21 (d9) | 11.8 |
|  | Ceftazidime | 2 (d5) | 1.5 (d7) | 2 (d7) | 1.8 |
| Ceftriaxone | Meropenem | 16 (d7) | 8 (d9) | 6 (d14) | 10 |
|  | Ciprofloxacin | 3.9 (d3) | 3.9 (d11) | 5.3 (d14) | 4.4 |
|  | Ceftazidime | 24 (d3) | 64 (d7) | 6 (d7) | 31.3 |
| Ertapenem | Meropenem | 8 (d9) | 31 (d7) | 48 (d9) | 29 |
|  | Ciprofloxacin | 1.5 (d9) | 2.6 (d7) | 2.6 (d11) | 2.2 |
|  | Ceftazidime | 1 (d1) | 1.5 (d5) | 3 (d11) | 1.8 |

MICs in the evolved populations remained appreciably higher than the control ones on Day 14 after incubating without any antibiotics for 72 hours. This suggests that the phenotypic changes induced by the NAPA persist despite removal of the antimicrobial pressure over this period.

Though the degree and timing of MIC changes varied between *P. aeruginosa* strains, the general patterns of change were consistent (as seen in Figure2). The increasing MIC became apparent following 1-3 days of incubation and usually peaked after 5-9 days. To relate these population-level resistance phenotypes to underlying genomic changes, we examined the temporal dynamics of resistance-associated mutations during exposure and across strains (Figure 2)

**Figure 2.**
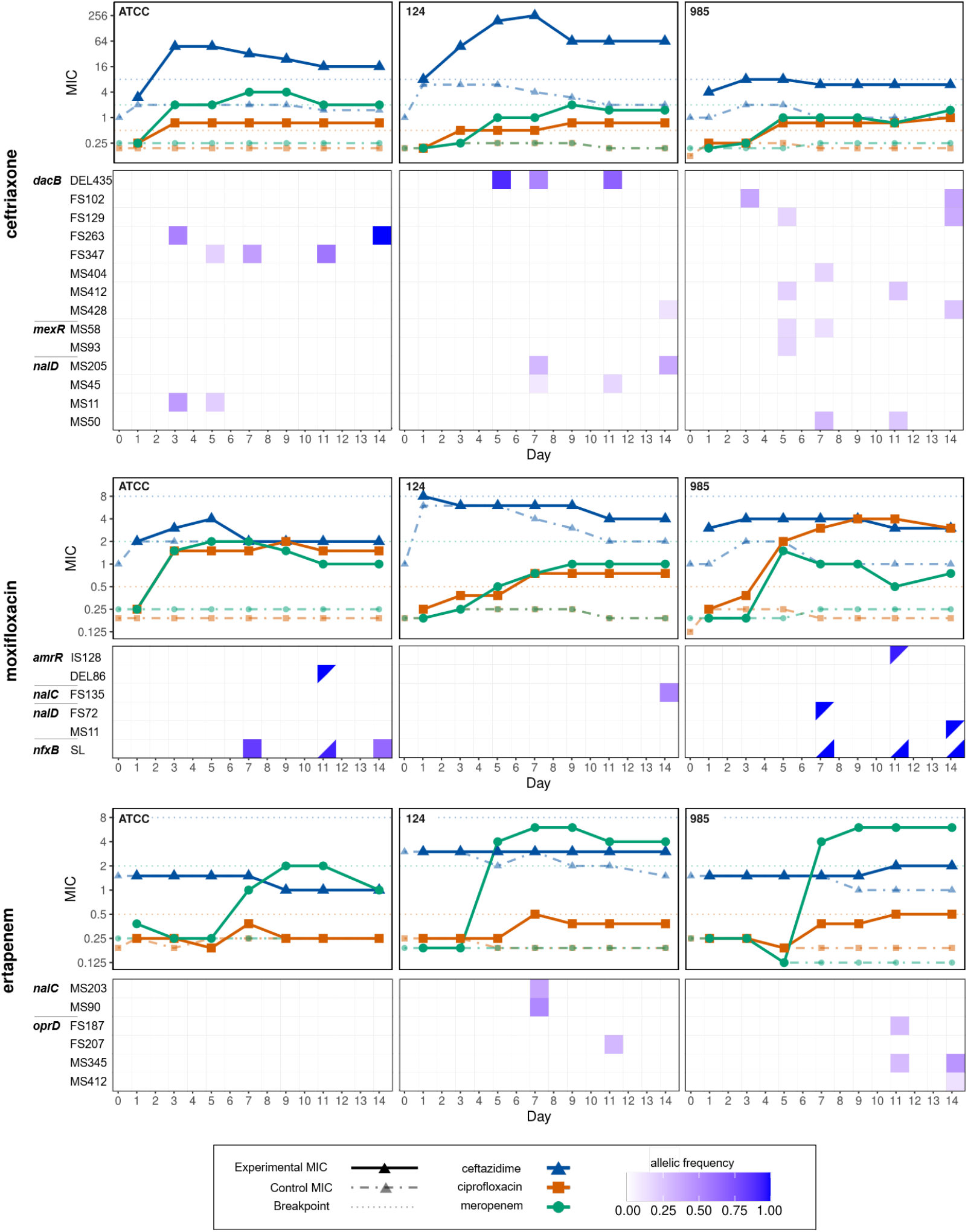
MIC trajectories and mutation allele frequency dynamics during serial exposure to non-antipseudomonal antibiotics. MIC trajectories and whole-genome sequencing results for all *Pseudomonas aeruginosa* strains and antibiotic exposures included in the study. For each strain–exposure combination, population-level MICs for antipseudomonal antibiotics are shown over time (upper panels), with corresponding allele frequency trajectories for resistance-associated mutations detected by whole-genome sequencing (lower panels). Allele frequencies reflect the proportion of sequencing reads supporting each variant at the indicated time point. Individual mutations are labeled by gene name. MIC measurements and genomic sampling were performed at discrete time points and are aligned temporally for visualization. The appearance, disappearance, and fluctuation of individual mutations highlight the heterogeneous and often transient nature of resistance-associated variants, while elevated MICs may persist despite turnover of specific alleles. No assumption of causality, fixation, or direct genotype–phenotype correspondence is implied, and alignment is intended to illustrate temporal coexistence rather than mechanistic linkage.

### Whole Genome Sequencing Results

Across all three strains, mutations emerged predominantly in genes regulating multidrug efflux systems (*nfxB, nalC, nalD, amrR*), porins (*oprD* family) and regulators of β-lactamase expression (*dacB*). A summary of selected relevant nonsynonymous single-nucleotide polymorphisms (SNPs) and small insertions/deletions and their allelic frequencies is presented in Table S3, and all recovered genomic changes in Tables S4-S6 Sample 86_784719_S63 (124 experiment, ertapenem exposure D3) generated a large number of SNPs relative to other isolates, and was removed from the analysis.

As illustrated in Figure 2, resistance-associated mutations emerged heterogeneously and transiently during periods of MIC escalation, with recurrent sampling of canonical resistance pathways rather than fixation of individual variants. The appearance of the mutations occurred during periods of MIC escalation (Figure 2) in several cases. Mutations in efflux pump repressors were among the earliest and most consistent findings. In moxifloxacin-exposed isolates, *nfxB* stop-loss and missense variants appeared by days 5–7, temporally coinciding with the steepest increases in ciprofloxacin MICs. These variants are predicted to derepress the MexCD-OprJ system, consistent with cross-resistance to fluoroquinolones and β-lactams. Similarly, different frameshift mutations in *nalC* and *nalD*—transcriptional repressors of MexAB-OprM—emerged in ceftriaxone- and ertapenem-exposed lineages in parallel across all three experiments, aligning with marked increases in ciprofloxacin and also ceftazidime and meropenem MICs.

Alterations in β-lactam–associated genes were also observed. Suspected loss-of-function mutations in *dacB* (encoding PBP4) occurred under both ceftriaxone and ertapenem exposure beginning on day 7. Disruption of *dacB* is a recognized driver of *ampC* derepression and consequent β-lactam resistance, providing a plausible explanation for the observed 16–48-fold increases in ceftazidime and meropenem MICs, possibly in combination with the mutations described above. *oprD* mutations were also observed in ertapenem-exposed isolates, and may possibly contribute to elevated meropenem MICs, as these molecules preferentially transport carbapenems.

Notably, resistance-associated variants were still detectable at day 14 in several lineages, paralleling the persistence of elevated MICs, although most mutations fluctuated in allelic frequency and were not fixed. Together, the sequencing results indicate that sub-MIC exposure to non-antipseudomonal antibiotics selects for heritable mutations that persisted for at least 3 days and that map to regulators known to derepress multidrug efflux systems and upregulate β- lactamase expression, consistent with broad collateral resistance phenotypes. Importantly, most SNPs and small SVs were not fixed and were identified intermittently, and at variable allelic frequencies over time (Table S3). Combining these findings with heterogeneity in colony morphology at different time points, this suggests the emergence of different subpopulations during exposure, which are selected out during MIC testing.

## Discussion

In this study, we demonstrate that prolonged exposure of *Pseudomonas aeruginosa* to subinhibitory concentrations of antibiotics lacking intrinsic antipseudomonal activity selects for stable, heritable resistance to antipseudomonal agents. Resistance emerged through convergent evolution and mutations in a set of regulatory genes governing multidrug efflux and β-lactamase expression—mechanisms classically associated with direct antipseudomonal pressure. The persistence of elevated MICs following antibiotic withdrawal indicates that these phenotypes reflect evolutionary adaptation rather than transient tolerance, revealing that canonical resistance pathways in *P. aeruginosa* can be selected independent of direct target engagement. The magnitude and persistence of MIC elevation further support stable evolutionary adaptation rather than transient tolerance. Whole-genome sequencing revealed parallel mutations across strains and antibiotic classes in efflux pump repressors (*nfxB, nalC, nalD, amrR, mexR*), a porin (*oprD*), and in *dacB*, a negative regulator of *ampC*. These changes are well-established mediators of multidrug resistance in *P. aeruginosa*. Derepression of efflux pumps through *nfxB* and *nalC/D* mutations likely upregulate the MexCD-OprJ and MexAB-OprM systems, enhancing export of fluoroquinolones and β-lactams^22^. Similarly, a loss of function in *dacB* under β-lactam exposure can derepress *ampC*, increasing resistance to meropenem and ceftazidime^23^. Although we focused on established AMR-related genes, we noted mutations in cell membrane and metabolic genes (for exmaple *sdhD, glgP*, see Tables S4-6) which may reflect metabolic and cell-envelope remodeling under chronic stress, that can enhance persistence and adaptive fitness, though may not reflect in the MIC^24–27^. Taken together, these convergent pathways suggest that subinhibitory exposure to antibiotics not targeting *Pseudomonas* directly can nevertheless trigger the same evolutionary routes exploited during direct antipseudomonal pressure.

The convergence on a limited set of regulatory nodes across multiple strains and antibiotic classes highlights the constrained evolutionary landscape through which *P. aeruginosa* rapidly adapts to antibiotic stress. In contrast to selection for novel resistance determinants, which may emerge under prolonged selection pressure particularly in the environment where horizontal gene transfers of exogenous AMR-determinants are more likely to occur, sublethal exposure to non-cognate antibiotics in culture repeatedly favored derepression of pre-existing multidrug resistance networks. This underscores the inherent plasticity and evolvability of the endogenous *P. aeruginosa* resistome.

Observational and experimental studies have identified antibiotic pressure as an important risk factor in the development of drug-resistant *P. aeruginosa*. A prospective cohort study in the ICU found that meropenem, ciprofloxacin, and ceftazidime administration was significantly associated with the development of drug-resistant PsA (adjusted HR 11.1, 4.1, 2.5 respectively)^28^. This association was further investigated *in vitro*. Feng et al.^29^ exposed a wild-type PsA strain to five different APA with stepwise increases in concentration over time. A subsequent gradual rise in MIC to each antibiotic was observed. Whole genome sequencing showed that the rise in ciprofloxacin MIC was linked to *gyrA, parC, gyrB* mutations, whereas that of meropenem and ceftazidime was related to increased beta-lactamase activity and not due to a consistently present genetic mutation^29^.

Like our study, Alyaseen et al’s directed evolution experiment showed that the ciprofloxacin-exposed PsA developed resistance to ciprofloxacin, but also cross-resistance to cefepime. This was likely mediated by upregulation of multidrug efflux pumps MexAB-OprM and MexCD-OprJ. The ciprofloxacin exposed strains did not however develop resistance to tobramycin, suggesting that MexXY-OprM was unaffected by the antimicrobial pressure^30^. Our results further confirm the importance of cross resistance but expand by demonstrating that the development can happen even under exposure to agents not traditionally considered to target *Pseudomonas*.

Fewer studies have examined whether the antimicrobial pressure of NAPA confers resistance to APA, and whether this association exists between different antibiotic classes. Thai et al.^31^ serially exposed a wild-type pan-susceptible PsA isolate to a steady low concentration of moxifloxacin. After 12 days of subculturing in moxifloxacin containing media, the resultant strain was highly drug resistant to moxifloxacin (MIC 1→ 128 mg/L), ciprofloxacin (0.0625→8 mg/L), and several other antibiotics. Genetic analysis revealed a deletion mutation (in *nfxB*) leading to increased expression of a multidrug efflux pump (MexCD-OprJ)^31^, similar to what we have found.

Together, these findings challenge the assumption that antibiotics lacking intrinsic antipseudomonal activity are evolutionarily neutral with respect to *P. aeruginosa*. Within the antibiotic classes examined, sublethal exposure was sufficient to select for canonical resistance architectures typically associated with direct antipseudomonal therapy. From an evolutionary perspective, these results suggest that resistance emergence in *P. aeruginosa* is less dependent on drug–target specificity than on the presence of sustained antimicrobial stress, with important implications for how antibiotic exposure histories are interpreted in both clinical and ecological contexts. In conclusion, our results suggest that within these classes, the concept of “*Pseudomonas* sparing” antibiotics does not, strictly speaking, exist within the antibiotic classes examined. Patients with *P. aeruginosa* infections should be considered to have elevated risks of resistance to anti-pseudomonal antibiotics if they have had recent exposure to non-pseudomonal antibiotics.

## Data Availability

Sequencing reads generated in this project will be available on NCBI SRA under BioProject accession PRNJA1419849. Kraken reports, reference sequences, annotations, variant call files (VCF) are available in the data supplement (10.5281/zenodo.18476895). Read alignments and filtered readsets are available from the corresponding author on reasonable request.

